# Cortical and subcortical substrates of minutes and days-long object value memory in humans

**DOI:** 10.1101/2023.03.17.533079

**Authors:** Sepideh Farmani, Kiomars Sharifi, Ali Ghazizadeh

## Abstract

Obtaining valuable objects motivates many of our daily decisions. However, the neural underpinnings of object processing based on human value memory are not yet fully understood. Here, we used whole-brain fMRI to examine activations due to value memory as participants passively viewed objects before, minutes after, and 1-70 days following value training. Significant value memory for objects was evident in the behavioral performance, which nevertheless faded over the days following training. Minutes after training, the occipital, ventral temporal, interparietal, and frontal areas showed strong value discrimination. Days after training, activation in the frontal, temporal, and occipital regions decreased, whereas the parietal areas showed sustained activation. In addition, days-long value responses emerged in certain subcortical regions, including the caudate, ventral striatum, and thalamus. Resting-state analysis revealed that these subcortical areas were functionally connected. Furthermore, the activation in the thalamo-striatal cluster was positively correlated with participants’ performance in days-long value memory. These findings shed light on the neural basis of value memory in humans with implications for object habit formation and cross-species comparisons.

## Introduction

Value-based memory, which relies on past stimulus-response associations, can evoke automatic responses independent of immediate outcomes (Dickinson & Balleine, 2002; Graybiel, 2008; Ashby et al., 2010; Ghazizadeh et al., 2016a). Such automatic behavior arises from stable object-value pairings formed through repeated experiences and can persist over a long period (Kim & Hikosaka, 2015; Anderson & Yantis, 2013). While habitual processes supported by value-based memory lead to efficient behavior on the positive side (Ghazizadeh et al., 2016b), on the negative side, they underlie disruptive attention to distractors (Anderson et al., 2011) and maladaptive behaviors in a number of neuropsychiatric disorders such as Tourette’s syndrome (Delorme et al., 2016) and drug addiction (Everitt & Robbins, 2005; Sjoerds et al., 2014).

A substantial body of work in animals has focused on the neural mechanisms of value-based memory. Electrophysiological evidence supports the role of the posterior basal ganglia in long-term value memory in non-human primates (NHPs) (Yasuda et al., 2012; Kim & Hikosaka, 2013; Kim et al., 2015). Furthermore, robust value discrimination of objects in regions of the temporal and prefrontal cortex, as well as in the striatum, amygdala, and claustrum, is demonstrated using fMRI (Ghazizadeh et al., 2018a). Notably, these differential responses persist for several months after training (Yasuda et al., 2012; Ghazizadeh et al., 2018a; Ghazizadeh & Hikosaka, 2021). The neural correlates of value memory in the human brain are remarkably less clear. A number of human studies aimed to distinguish neural representations of goal-directed and habitual processing across the whole brain, suggesting that the dorsolateral striatum is involved in habitual processing, whereas the dorsomedial striatum and orbitofrontal cortex (OFC) are implicated in goal-directed processing (Tricomi et al., 2009; de Wit et al., 2012; Wunderlich et al., 2012; Graybiel & Grafton, 2015; Glascher et al., 2010; Liljeholm et al., 2012; Daw & O’Doherty, 2014; McNamee et al., 2015). Moreover, previously reward-associated objects, despite being task-irrelevant, are shown to evoke preferential responses in the caudate tail, intraparietal sulcus (IPS), and extra-striate visual cortex (Anderson et al., 2014). In addition, the ventral striatum (VS) has recently been shown to be involved in retaining long-term value memory of objects (Kang et al., 2021). However, the whole-brain functional circuitry of object value memory spanning minutes to days remains unexplored.

In the current study, participants were trained to associate novel fractal stimuli with monetary reward or no reward. Brain responses to the passive viewing of objects were measured before, minutes after, and days after the final training session using fMRI. Value training was conducted across three sessions to elicit stable value-based memory (Wimmer et al., 2018). We adopted the passive-viewing paradigm previously used to investigate the long-term value memory of objects in NHPs (Ghazizadeh et al., 2018a; Ghazizadeh et al., 2018b), which allowed us to explicitly examine the modulations in the visual processing of objects attributed to value memory, from minutes to days. Task and resting state fMRI results revealed a network of parietal, occipital, and frontal cortical regions demonstrating value memory immediately after training, while in the days-long memory, the thalamo-striatal areas showed significant activations.

## Results

Participants learned the value of 40 fractal objects randomly associated with a monetary reward (good objects) or no reward (bad objects) across three sessions (Fig. 1A). The large number of stimuli and arbitrary value assignments resembled real-life experiences involving many objects and ensured that the contrasts between good and bad objects were not confounded by the idiosyncratic physical features of objects (Fig. 1B). During the training, after a central fixation offset, a single fractal was displayed on the left or right side of the screen. Participants had to select that object by pressing the left or right key, and then the associated reward was presented (80% force trials). Interspersed among force trials were unitary choice trials in which single objects were presented, and the participants were instructed to indicate the value of the objects (20% choice trials) (Fig. 1C). No monetary feedback was provided in the unitary choice trials. Following the value training sessions, the neural responses elicited by passive exposure to good and bad objects were measured using fMRI, minutes (post-training fMRI session for minutes-long memory), and within 1-70 days after the third value training session (memory fMRI session for days-long memory) in a block design paradigm consisting of base blocks with only central fixation and probe blocks with a single fractal flashing in the center or periphery (Fig. 1E) (see methods for details). As a control, the same passive-viewing fMRI paradigm was performed prior to the first training session (pre-training fMRI session, Fig. 1A). Participants were instructed to keep the central fixation during the passive-viewing task, and a fixation-break classification indicated no significant difference in the number of fixation breaks between good and bad objects (t12=0.04, p=0.96) (see methods).

**Fig. 1.**
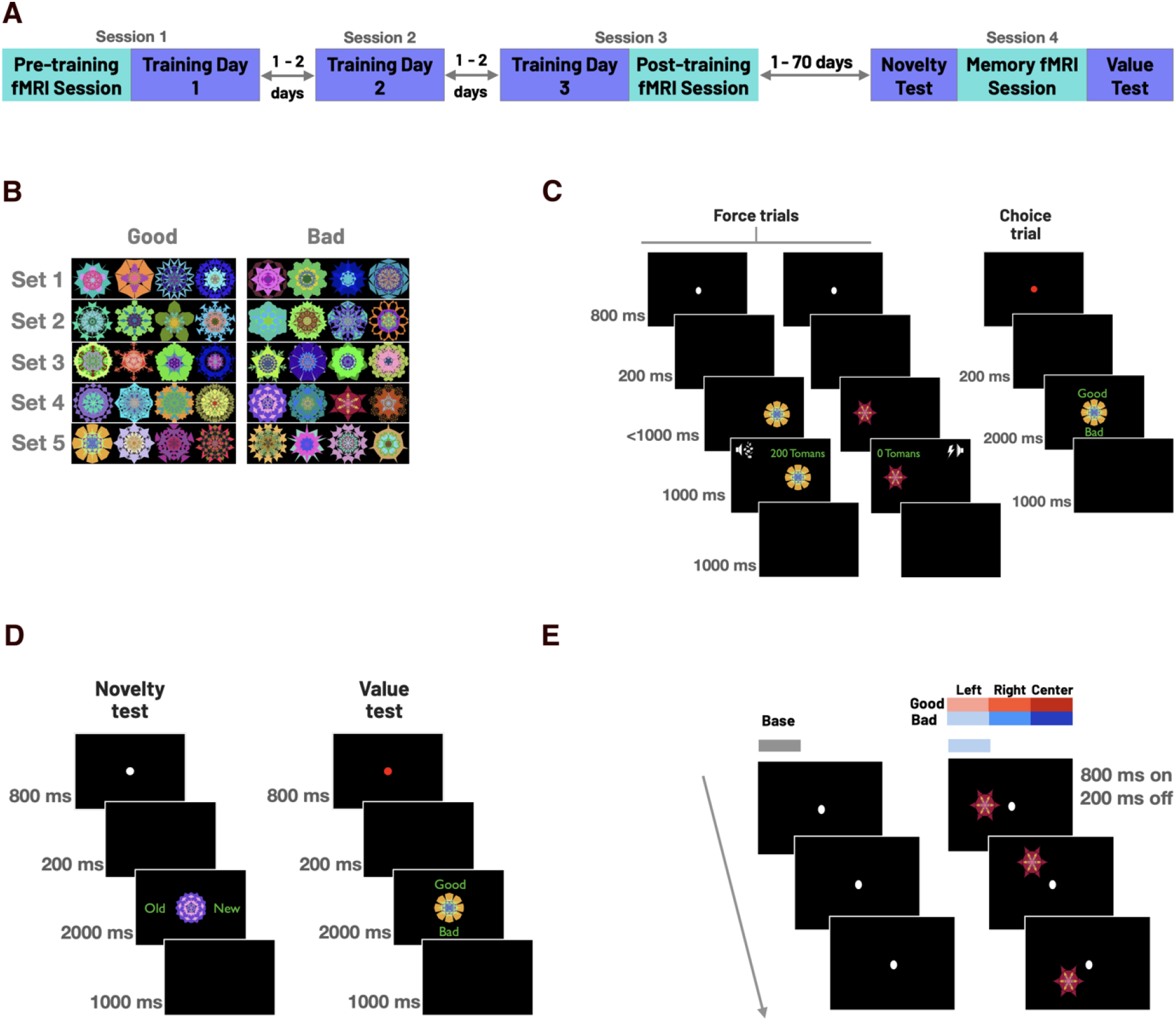
Experimental timeline and task paradigms. (A) Experimental timeline: Participants learned the value of fractal objects across three sessions. Brain responses were measured using fMRI before, immediately after, and within 1-70 days after the last training session. (B) Stimuli: 40 fractals were randomly assigned into two groups of good and bad objects (the assignment switched across subjects). (C) Value learning task consisted of two types of trials. In force trials, fractal objects were repeatedly associated with a monetary reward (good objects) or no reward (bad objects). In choice trials, participants were instructed to choose the value of a single object shown in the center by pressing the up or down keys without receiving any feedback about the accuracy or monetary outcome. (D) In the novelty test, participants were instructed to indicate the familiarity of a single object shown in the center. The value test was the same as choice trials in the training sessions. (E) fMRI Passive-viewing task consisted of alternating base and probe blocks. In the base blocks, participants looked at the central fixation, and no object was shown. In the probe blocks, a single good or bad object was shown in the center, on the left, or the right hemifield.

A repeated-measures ANOVA performed on the accuracy in choice trials yielded a significant improvement by training (F2,120 = 11.99, p < 0.001), a significant effect of value (F1,120 = 14.87, p = 0.002), and no significant interaction (F2,120 = 0.0035, p = 0.75) (Fig. 2B). Performance significantly increased across the three training sessions (session 1: M = 0.86, SD = 0.01; session 2: M = 0.90, SD = 0.01 and session 3: M = 0.94, SD = 0.009). Furthermore, participants showed higher accuracy for good vs. bad objects in the last two sessions (session 2: t20= 2.8, p = 0.01; session 3: t20= 3.25, p = 0.004). In force trials, performance was high throughout the training sessions (M > 0.99, SD < 0.003) (Fig. S1). In addition, an analogous ANOVA performed on the choice response time showed a significant effect of training with participants becoming faster (F2,120 = 20.48, p < 0.001), but no effect of value (F1,120 = 3.67, p = 0.06) nor interaction (F2,120 = 0.15, p = 0.86) (Fig 2C). Response time significantly decreased across the three training sessions (session 1: M = 1.1, SD = 0.03; session 2: M = 0.99, SD = 0.03 and session 3: M = 0.9, SD = 0.02). Moreover, in the final training session, participants responded faster to good objects compared to bad objects (t15 = -2.78, p = 0.01), consistent with previous literature showing faster response to higher value objects (Wimmer et al., 2018; Tankelvitch et al., 2020; Kim et al., 2015). The reduction in response time was also observed in force trials (Fig. S1). The differential responses to good and bad objects, and ∼90% choice accuracy for object values, demonstrated that participants successfully learned the value associations across the training sessions.

**Fig. 2.**
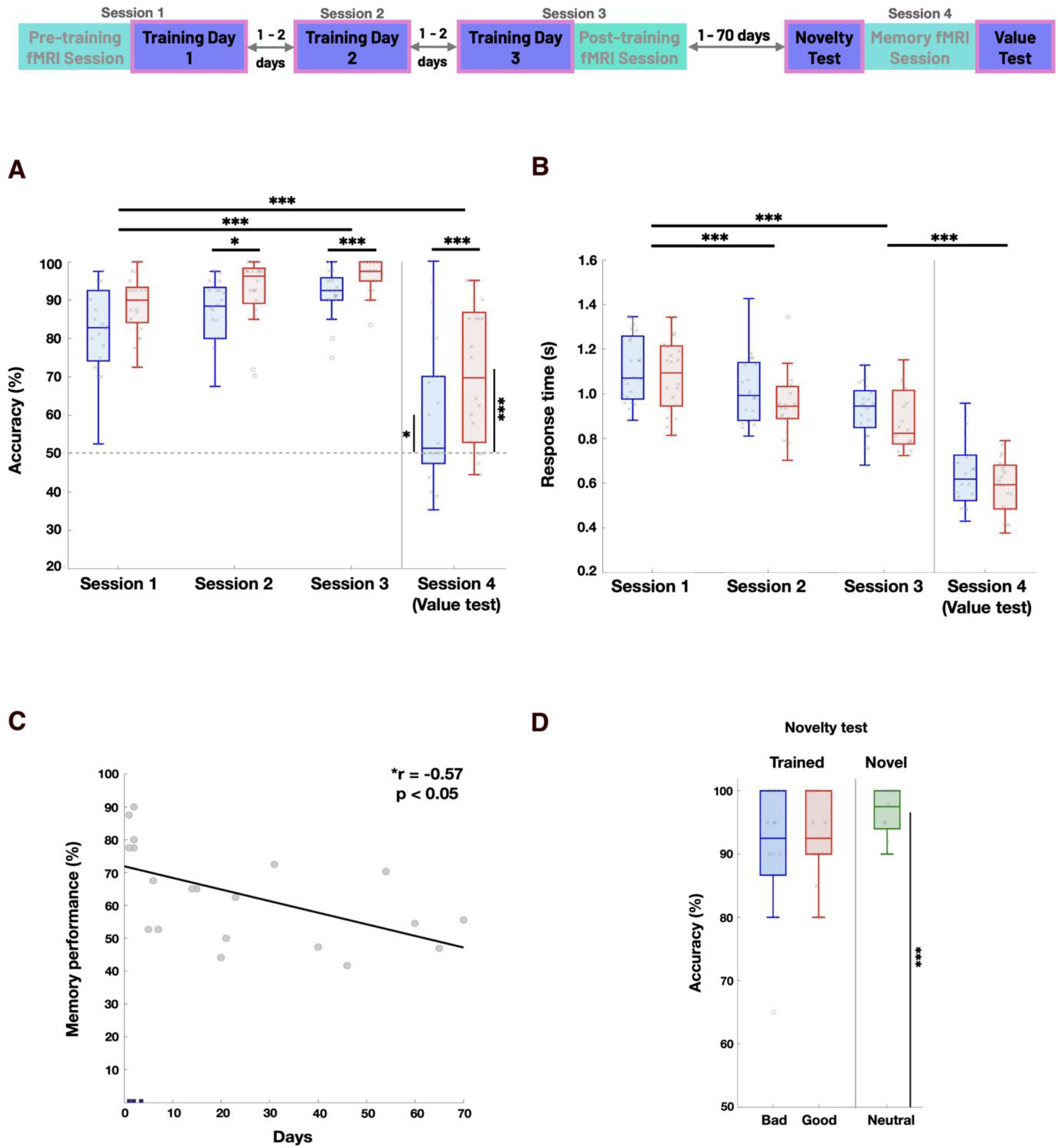
Humans can readily learn and form days-long value memory of a large number of abstract fractal objects. (A) Behavioral results are from choice trials in the training sessions highlighted in light pink. (B) Choice performance across the three training sessions and the memory session. Boxplots show the choice rate of good and bad objects in each session. The central mark within each box indicates the median, and the bottom and top of the box correspond to the 25th and 75th percentiles, respectively. The whiskers extend to the extreme values within 1.5 times the interquartile range. Outliers are plotted as dots. (C) Response time in choice trials across the three training sessions and the value test. Same conventions as in (B). (*p < 0.05, **p < 0.01, ***p < 0.005) (D) Choice performance in the value test vs. the duration of the intervening period from the last training session (Pearson’s correlation “r” and significance “p” are noted in the plot). (E) Choice performance in the novelty test for good, bad, and novel stimuli. Same conventions as in (B).

A days-long value test was performed 1-70 days after the final training session. Despite a significant decrease in average performance for both good and bad objects compared to the initial training session (t20 = -5.19, p < 0.005), performance was still significantly above chance for both object types (t15 = 2.55, p = 0.02 for bad and t20 = 5.41, p < 0.001 for good objects), with performance for good objects remaining higher than bad objects (t20 = 3.58, p = 0.001) (Fig 2B). Choice response time for good and bad objects in the value test further decreased compared to the final training session (t20 = -6.93, p < 0.001) (Fig 2C). As expected, the decline in value memory was significantly and negatively correlated with time since the last training session (Fig. 2D).

Reduced performance in session 4 (value test) could have been caused by forgetting the object values or the objects themselves. In the first scenario, participants should have remembered that they had seen an object before but should have been skeptical about its value, while in the second, the object should have looked novel to them. To address this possibility, some of our participants took part in a novelty test before the memory scan, in which good and bad objects were presented along with the same number of novel objects. Participants were instructed to choose whether the object was novel or familiar. Notably, the average performance for old and new objects was 92% and 96% (p < 0.001), respectively, with no difference between good and bad objects (t12 = -1.85, p = 0.09) (Fig. 2E). These results suggest that participants had a clear idea of which objects were novel and that the errors in value judgments were not due to forgetting previously seen objects, but most likely to forget their value.

### Brain areas involved in object value memory

To evaluate the neural substrates of value-based memory, we employed a passive-viewing task with no reward delivery. Good and bad objects were presented in the scanner in the center, or the right or left hemifield, and participants were instructed to simply keep their eyes on the central fixation point (Fig. S2). Given the large number of objects, their random visual features, and their arbitrary assignment to good and bad groups, differential responses to good and bad objects could only arise due to previously learned values. As expected, we found no significant difference in responses to good and bad objects before training (pre-learning session, p < 0.05, alpha = 0.01, cluster corrected). In contrast, significant differentiations were observed in several brain regions minutes after the last training session (post-training fMRI session) in the intraparietal sulcus (IPS), middle and inferior occipital gyrus, insula, fusiform gyrus (FFG), and dorsolateral putamen (Fig. 3B, Table 1, Fig. S3). These regions were more strongly activated in response to good objects than bad ones. Limiting the good vs. bad contrast in the post-learning session to objects correctly identified in the final training session (memorized objects) yielded similar results due to the small number of errors.

**Fig. 3.**
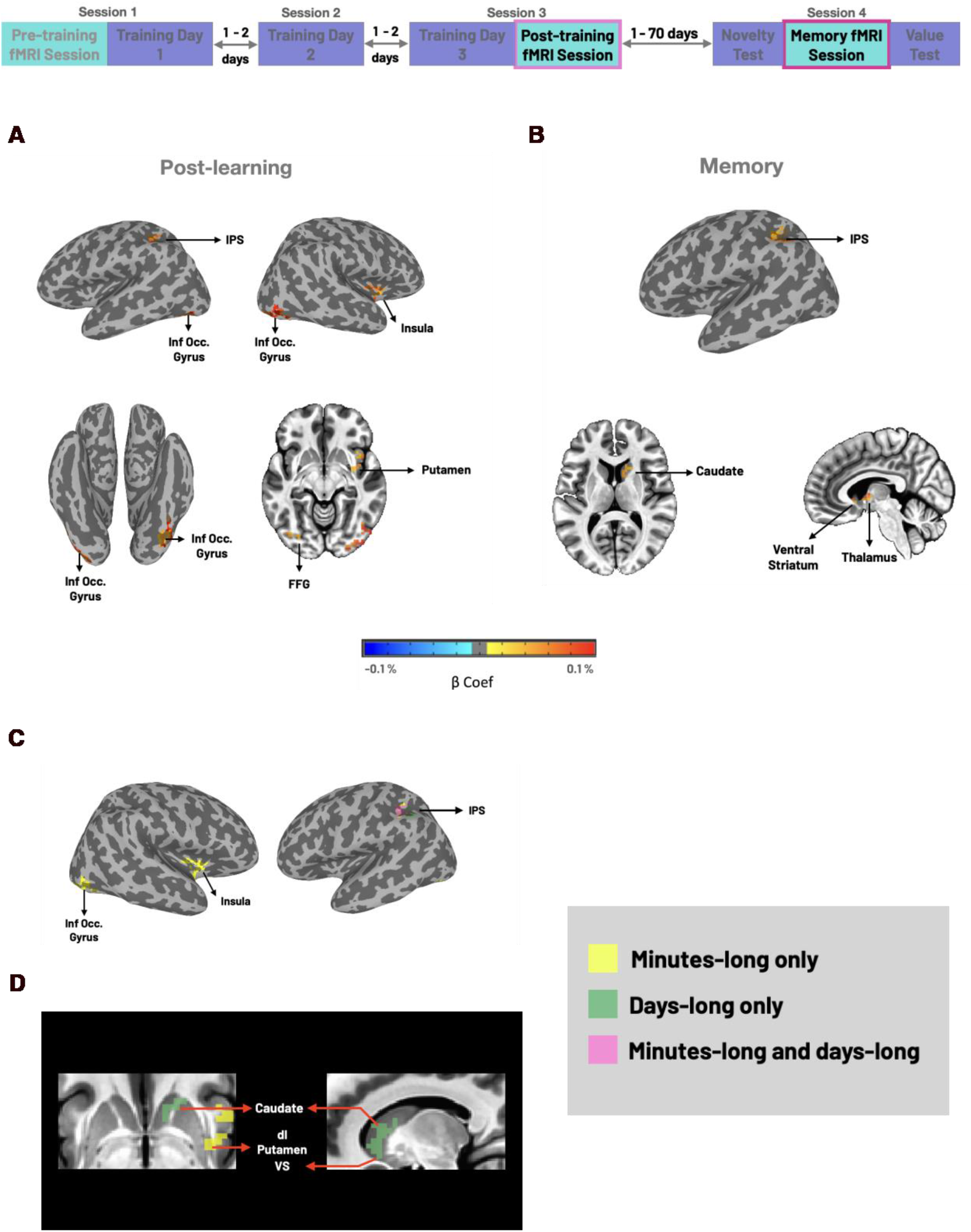
Whole-brain neural correlates of minutes-long and days-long value memory. (A) Experimental timeline: post-learning and memory fMRI sessions are highlighted in light and dark pink, respectively. (B) Brain regions with significant differential responses to good and bad objects in the post-learning session (minutes-long memory) (p < 0.05, α < 0.05, cluster corrected). (C) Same as B but for the memory fMRI session (days-long memory) (p < 0.05, α < 0.05, cluster corrected). (D) Cortical regions with significant good vs. bad discrimination in the post-learning session (minutes-long, yellow), in the memory session (days-long, green), and both sessions (pink). (E) Same as D but for subcortical regions.

**Table 1.** Summary of whole-brain analysis results.

| **Contrast** | **Regions** | **Cluster size** | **x** | **y** | **z** |
| --- | --- | --- | --- | --- | --- |
| **Good vs. bad post-learning** | R insular lobe | 72 | 43 | 10 | -5 |
|  | R dorsolateral putamen |  |  |  |  |
|  | R inferior occipital gyrus | 62 | 40 | -93 | -9 |
|  | R middle occipital gyrus |  |  |  |  |
|  | L intraparietal sulcus | 49 | -41 | -44 | 47 |
|  | L inferior parietal lobule |  |  |  |  |
|  | L superior parietal lobule |  |  |  |  |
|  | L fusiform gyrus | 47 | -35 | -84 | -12 |
|  | L inferior occipital cortex |  |  |  |  |
| **Good vs. bad memory** | R ventral striatum | 82 | 10 | 7 | -6 |
|  | R dorsal caudate |  |  |  |  |
|  | R thalamus |  |  |  |  |
|  | R ventral caudate |  |  |  |  |
|  | L intraparietal sulcus | 70 | -43 | -50 | 47 |
|  | L inferior parietal lobule |  |  |  |  |

Clusters of activity exceeding whole-brain $p < 0.05$ , $\alpha < 0.01$ , cluster-corrected (cluster size = 46 voxels for the post-learning and cluster size = 60 voxels for the memory fMRI session).

Object value discrimination based on days-long memory was found in a subset of regions activated in the post-training session, including the intraparietal and inferior occipital gyrus. (Fig. 3C, Table 1, Fig. S3). In contrast, certain areas, including the FFG, insula, and dorsolateral putamen, did not pass the cluster correction in the memory session. Similar results were observed when limiting the good vs. bad contrast to memorized objects in the value test (Fig S4, Table S1). Interestingly, robust value coding was observed in several subcortical areas in the memory fMRI session that were not significantly activated in the post-training scan. This value coding was highly evident in the caudate, thalamus, and ventral striatum (VS) (Fig. 3C). Together, these results suggest that activity in the cortical areas was more pronounced in minutes-long memory. In contrast, certain subcortical regions, such as the basal ganglia and thalamic circuits, demonstrated enhanced activation in days-long memory (Fig 3D-E).

### Resting-state networks are differentially engaged in maintaining object value memory

The appearance and disappearance of certain cortical and subcortical areas in post-learning and memory sessions suggested that the observed activations might be organized at the network level. To investigate the network properties of value-based memory, we leveraged the functional connectivity among value-coding regions during rest. Pairwise correlations between the resting-state time course of regions significantly active in either the post-training or memory session were computed and converted into a distance matrix (see methods). A 2D embedding of areal distances was implemented using multidimensional scaling, and areas were clustered using a density-based clustering algorithm (Ghazizadeh et al., 2018a). In this embedding, each point corresponded to one area, and a smaller distance between the two points indicated a stronger resting state correlation (Fig. 4A). The results demonstrated that the value-coding regions could be divided into five clusters based on functional connectivity, namely: 1. Occipito-temporal cluster, 2. Parietal cluster, 3. Fronto-striatal cluster, and 4. Thalamo-striatal cluster. We compared the average value signal across each cluster in the post-training and memory sessions to gain further insight into the relationship between these clusters and the value signal. The results indicated that in the thalamo-striatal cluster, the value signal was significantly higher in the memory session than in the post-training session, while the opposite effect was observed in the occipito-temporal and fronto-striatal clusters (Fig. 4B). Furthermore, the value signal in the post-learning session showed a significant decrease with the distance from the center of mass of the occipito-temporal and fronto-striatal clusters in the functional space, whereas the value signal in the memory session was positively correlated with the functional distance from the center of the same clusters. Conversely, the value signal in the post-learning increased significantly with the functional distance from the center of mass of the thalamo-striatal cluster, while the value signal in the memory session showed a decreasing trend with the functional distance from the center of mass of this cluster. Together, these results showed that areas functionally connected to the occipito-temporal and fronto-striatal clusters vs. the thalamo-striatal cluster played an essential role in encoding minutes-long vs. days-long value memory (Fig. 4 C-D).

**Fig. 4.**
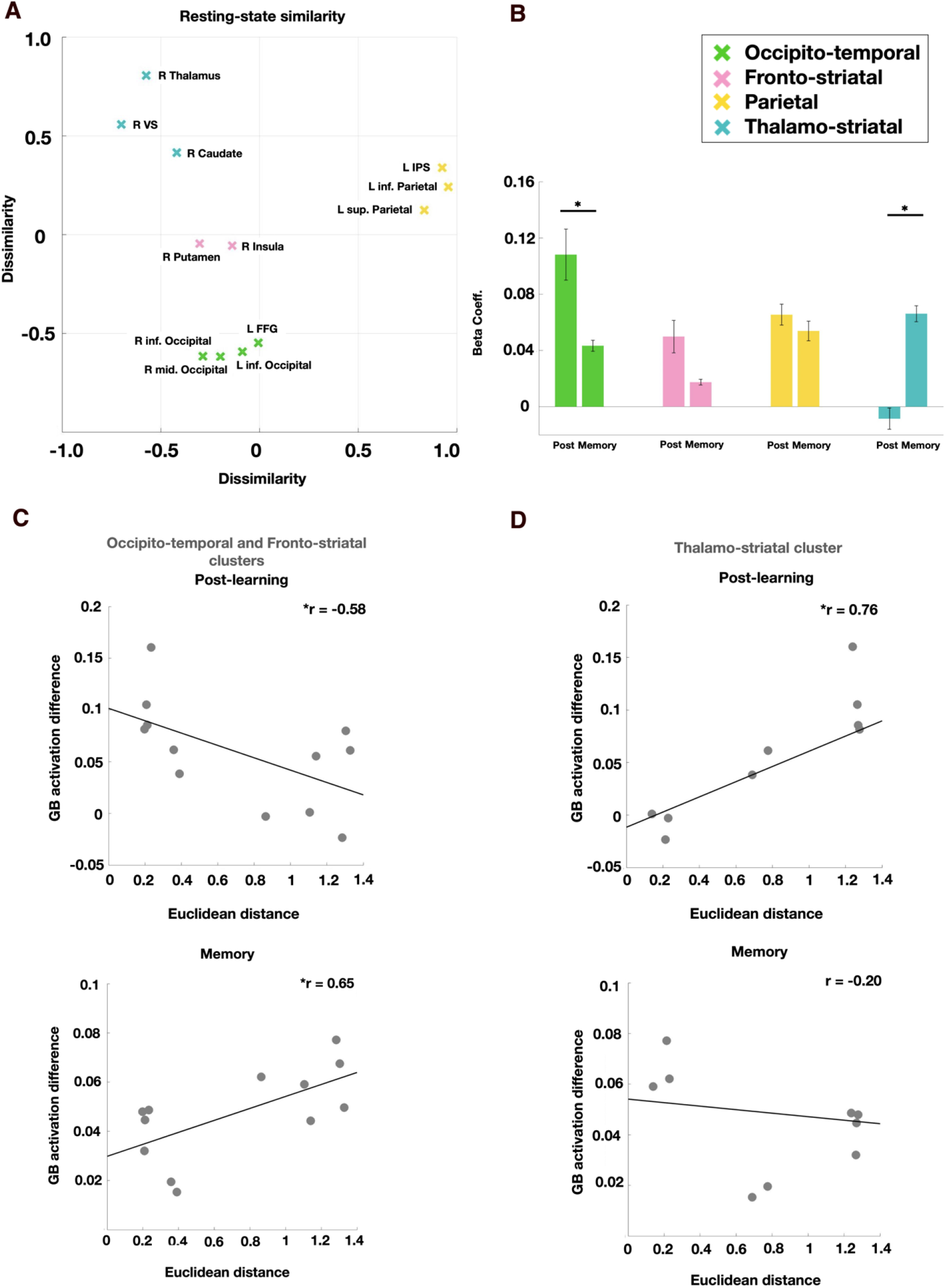
Functional connectivity between regions activated by minutes- or days-long value memory. (A) Functional distance of brain regions with significant value discrimination shown in 2D space using low dimensional embedding and clustered using a density-based algorithm. Clusters are plotted with different colors. (B) Good vs. bad beta coefficients for different clusters in post-learning and memory fMRI sessions. (C) Good vs. bad activation in post-learning and memory sessions as a function of the Euclidean distance from the center of mass of the occipito-temporal and fronto-striatal clusters. (D) Same as C but for the Euclidian distance from the center of mass of the thalamo-striatal cluster. (Pearson’s correlation “r” is noted in (C), (D), *p < 0.05, **p < 0.01, ***p < 0.005).

### Activation in the thalamo-striatal cluster correlates with days-long memory performance

Our results revealed regions with significant value discrimination in the memory session. Among these regions, activation in the thalamo-striatal cluster, including the right caudate nucleus, right thalamus, and right ventral striatum passed the cluster threshold for the memory > post-learning contrast (Fig. 5C). However, it is not clear whether these areas or other clusters contributed to the participant’s behavior in the value test. Remarkably, the magnitude of value-dependent activation in the thalamo-striatal cluster showed a significant positive correlation with performance in the value memory session (Fig. 5A, Fig. S5). In our experiment, days-long value memory was tested over a relatively long period of 1-70 days. As shown previously, memory performance was dependent on the time elapsed since the last training session and decreased significantly over time (Fig. 1D). Similar to performance, activation in the thalamo-striatal cluster decreased significantly with the duration of the intervening period (Fig. 5B, Fig. S5).

**Fig. 5.**
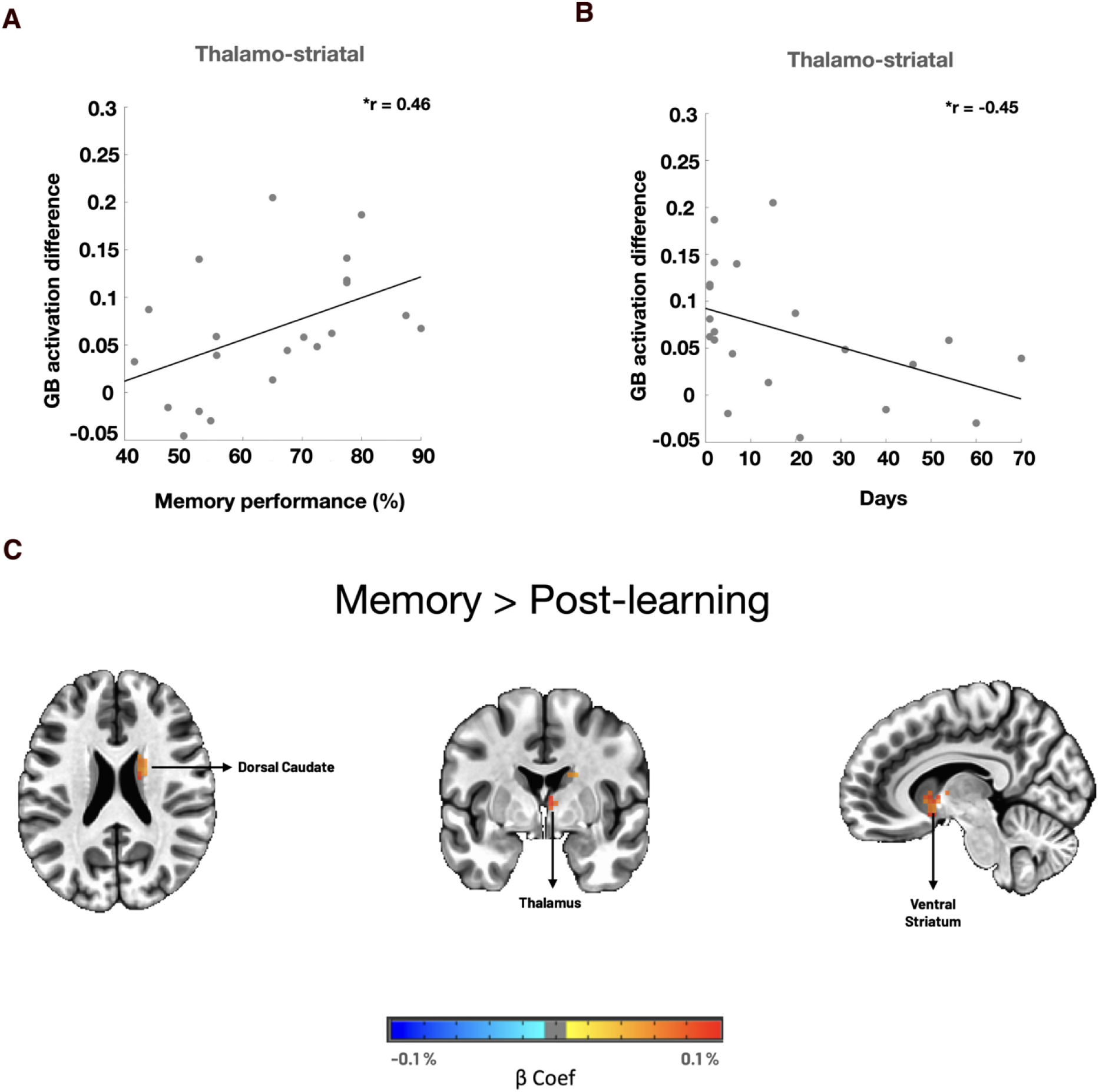
Activation in the thalamo-striatal cluster correlates with days-long value memory performance. (A) Good vs. bad beta coefficient averaged over the thalamo-striatal cluster as a function of participants’ performance in the memory session. (B) Same format as (A) but as a function of time elapsed since the last training session (Pearson’s correlation “r” is noted in (A) and (B)). *p < 0.05, **p < 0.01, ***p < 0.005). (C) Areas activated by the memory minus post-learning value contrast (p < 0.05, α < 0.05, cluster corrected, cluster size = 59 voxels).

## Discussion

Building on previous animal research that has demonstrated robust sensitivity to object value memory in multiple cortical and subcortical areas (Ghazizadeh et al., 2018a), we investigated the neural underpinnings of value-based memory in humans. Participants were trained across three sessions to learn the value of novel fractal objects. Neural responses were measured using fMRI before, 15-30 minutes after (minutes-long memory), and days after (days-long memory) the last training session. We employed a passive-viewing task in the absence of reward to assess visual responses modulated by the previously learned values of objects. Minutes after the final training session, we found enhanced responses in the middle and inferior occipital gyrus, FFG, insula, and IPS (Fig. 3). Despite the time-dependent decline in performance, participants were able to remember the value of many objects significantly above chance days after the last training (Fig. 2). This decline in performance was specific to forgetting the object values, not the objects themselves (Fig. 2E). Imaging results during this time showed persistent activation in some areas, including the IPS. Remarkably, we observed the emergence of value coding in the caudate, thalamus, and VS in the memory session (Fig. 3). Further investigation revealed that the activated regions in either scan formed several functional networks with different behaviors in minutes-long and days-long memory (Fig. 4). While the functional distance from the occipital and parietal clusters determined the strength of minutes-long value memory, it was the distance from the thalamo-striatal cluster that determined the strength of days-long memory. Among the regions with strong days-long value memory, the subcortical regions showed greater activation in the memory vs. post-learning scans, which was significantly correlated with participants’ performance in the days-long value test (Fig. 5).

Minutes after training, regions in the occipito-temporal, parietal, and fronto-striatal clusters exhibited enhanced value coding. Value discrimination in the insula, temporal, and occipital areas is in accordance with previous studies showing value-based modulation of representations in the cortical regions (Anderson, 2016; Qi et al., 2013; Hickey and Peelen, 2015; Tankelevitch et al., 2020; Ataei et al., 2022). Nevertheless, activation in several cortical areas decreased significantly across days. One possible explanation could be that some of these areas are more involved in the minutes-long value memory of objects, and as consolidation takes place, memory is stored elsewhere in the brain. Among these fading areas, the temporal and occipital cortices may be involved in regulating and forming lasting value memories but could disengage once the memories are ingrained (Sligte et al., 2009). On the other hand, the IPS showed persistent value responses days after training, consistent with previous findings, indicating the modulated representations of previously reward-associated stimuli in the IPS (Anderson et al., 2014; Ghazizadeh et al., 2018a; Platt et al., 1999; Peck et al., 2009). Our results extend these previous findings and implicate IPS in the long-term consolidation of reward memory in humans.

It is known that repeated pairing of objects with rewards induces habitual processing and leads to the formation of stable value memories (Kim & Hikosaka, 2015; Kim et al., 2015; Yasuda et al., 2012; Ghazizadeh et al., 2016b). Previous studies have provided evidence for the involvement of the dorsolateral striatum in habitual processing (Tricomi et al., 2009; de Wit et al., 2012; Wunderlich et al., 2012; Lee et al., 2013). These findings accord with electrophysiological studies in animals (Yin & Knowlton, 2006; Kim & Hikosaka, 2013). Consistent with these results, we found pronounced activation in the dorsal and ventral striatum extending to the thalamus days after training, which was significantly correlated with participants’ behavioral days-long memory (Fig. 3,5). In a previous study, the VS has been shown to be significantly activated in response to high-value objects presented centrally 2-5 days after value training (Kang et al., 2021). This aligns with our results, demonstrating the emergence of activity in the VS in the days-long memory session. However, in contrast to our results, in the aforementioned study, no activation was found beyond the VS in other cortical and subcortical areas, and also no behavioral measure of participants’ performance in the days-long memory session was provided. One possibility for the discrepancy between the whole-brain results is the differences in task designs. In Kang et al. (2021), object values were presented via binary choice trials, which could lead to a selection bias, modulating brain responses irrelevant to reward value (Failing & Theeuwes, 2018). In our task, selection bias was avoided by training participants using force trials and probing the value learning using unitary choice trials (Fig. 1). In addition, in our passive-viewing task, objects were presented in both the periphery and the center, which might have resulted in the engagement of several cortical and subcortical areas that are known to have lateralized receptive fields (Hasson et al., 2002; Corbetta et al., 2002).

Extensive work in human literature has focused on value-driven attentional capture (VDAC), which has revealed the influence of value learning on attention. This influence persists even when the previously reward-associated stimuli are currently task-irrelevant (Della Libera & Chelazzi, 2006; Hickey et al., 2010; Anderson et al., 2011; Anderson & Yantis, 2013; Anderson, 2016; Failing & Theeuwes, 2018). Neuroimaging studies have shown that the VDAC leads to greater activity in the visual cortex (Saproo & Serences, 2010; Hopf et al., 2015; MacLean & Giesbrecht, 2015; Anderson et al., 2014; Hickey & Peelen, 2015; Anderson, 2017), parietal cortex (Anderson et al., 2014; Barbaro et al., 2017; Lee & Shomstein, 2013; Qi et al., 2013; Anderson, 2017), and the caudate tail (Anderson et al., 2014; Anderson, 2016; Moore & Zirnsak, 2017). Brain responses contributing to salience coding (such as VDAC) and value-based memory are highly entangled. Therefore, some of the activated areas in the current study may also be involved in other processes. Indeed, it is known that the insula and thalamus encode stimulus salience irrespective of the valance (Seeley, 2019; Zhou et al., 2021). Nevertheless, we emphasize that regardless of the interpretation and downstream functionality of the observed activations in our study, they were all caused by the value memory of objects. Future studies are needed to elucidate the distinct functionality of neural representations attributed to attentional, emotional, motivational, or pure valuation domains.

In summary, our results revealed cortical and subcortical networks spanning minutes-long to days-long memory of previously learned object values. While the occipito-temporal, fronto-striatal, and parietal networks were activated within minutes after value learning, the parietal areas and the thalamo-striatal functional circuitry retained the value memory for days. Although the habitual mechanisms underlying value-based memory can render efficient visual processing of objects (Ghazizadeh et al., 2016b) and free up precious cognitive resources for new experiences, the role of habit formation in maladaptive behaviors cannot be overstated (Delorme et al., 2016; Everitt & Robbins, 2005; Sjoerds et al., 2014). Causal manipulation in these areas using electrical, magnetic, chemical, or genetic methods to disrupt such value-based maladaptive behaviors is a crucial direction for future investigations (Luigjes et al., 2019; Song et al., 2019; Antonelli et al., 2021).

## Materials and Methods

### Participants

26 healthy adults with normal or corrected-to-normal vision participated in the present study. Five participants were excluded due to either incomplete sessions or excessive head motion. Our final sample included 21 participants (10 females, mean age: 22.23 years, range:19–32). All participants provided written informed consent and were compensated for their participation with a flat monetary reward and an additional monetary bonus based on their performance on the value learning task. The experimental protocol was approved by the ethics committee of the Institute for Research in Fundamental Sciences (IPM) (protocol number 98/60.1/3025).

By performing power analysis on data from a pilot experiment (n = 6, mean beta = 0.02, SD = 0.019), we found that the estimated sample size to achieve an 80% power (β=0.2) at α=0.05 for detecting a difference between the good and bad objects in the caudate was 10 (Faul et al., 2009).

### Stimuli

80 fractal images similar to those used to study the neural basis of value-based memory in NHPs were utilized (Ghazizadeh et al., 2018a) (Fig. 1B). The parameters, including size, color, etc., were randomly selected for each fractal. 40 fractals were used in the value learning task. Half of these stimuli were associated with reward (good objects), and the other half received no reward (bad objects) during training. The order of objects was counterbalanced across participants. The remaining 40 fractals were used as novel stimuli in the novelty test.

### Experimental Procedure

To examine the neural substrates of value-based memory, participants were trained over the course of three sessions outside of the scanner, and their brain responses were measured across three rounds of fMRI scanning. The scans took place before the initial training session (pre-learning session), immediately after the final training session (post-learning session), and one or more days (1-70) after completing the training (memory session) (Fig. 1A). All tasks were presented using MATLAB (MathWorks) and Psychtoolbox-3 (Kleiner et al., 2007).

### Value Learning Task

Participants completed three training sessions, each consisting of five blocks. Each block was performed with a set of 8 fractals (4 good and 4 bad fractals) and comprised 80 trials, 64 force trials, and 16 choice trials. On each force trial, a white fixation dot was presented for 800 ms, followed by a blank screen for 200 ms (Fig. 1C). An object was then presented on the left or right side of the screen for 1000 ms, and participants were required to indicate their choice by pressing the left or right button. After the selection, visual and auditory feedback indicated the outcome for 1000 ms. The successful performance led to a reward (+200 tomans) for the good objects and no reward (0 tomans) for the bad objects. Each trial was followed by an intertrial interval (ITI) of 1000 ms. Choice trials were interspersed among the force trials, yielding training blocks consisting of 4 force trials and 1 choice trial. Each choice trial started with the presentation of a red fixation dot for 800 ms, followed by a blank screen for 200 ms. An object was then presented for 2000 ms in the center, along with the options “good” and “bad” above or below the object (the position of the options was randomized across subjects). Each trial was followed by an intertrial interval (ITI) of 1000 ms. Participants were instructed to indicate the object’s value by pressing the up and down arrow keys on the keyboard. No feedback was given on the accuracy of the choice.

### Novelty Task

12 participants underwent a novelty test before the memory scan. The task comprised 80 trials. Each trial started with the presentation of a fixation dot for 800 ms, followed by a 200 ms blank screen (Fig. 1D). An object was then presented in the center for 1000 ms. Participants were required to choose whether they had seen the objects in the training sessions (old) or not (new) by pressing the left and right arrow keys on the keyboard. No feedback was given throughout the task. Each trial was followed by an ITI of 2000 ms. The order of the objects was counterbalanced across participants.

### Passive-viewing Task

Neural responses to good and bad objects were measured before the first training session, immediately after, and within 1-70 days of the final training session. Participants engaged in a passive-viewing task across 7 runs. Each run consisted of 36 blocks (18 base and 18 probe) of 10 s duration (Fig. 1E). There were 6 different probe blocks: [good, bad] x [left, center, right]. The order of the blocks was pseudorandomized, and each stimulus was presented three times during each session, once in each of the left, center, or right hemifields. During each probe block, a fractal was flashed (800 ms on, 200 ms off) in the center or one hemifield (∼6^◦^ eccentricity across horizontal and 45^◦^ oblique directions) while the participant maintained central fixation. The base blocks had the same duration and contingencies as the probe blocks, except that no object was shown, and participants only had to keep their eyes on the central fixation. The order of objects was pseudorandomized across participants.

### Resting-state fMRI

In addition to the passive-viewing task, 18 participants completed resting scans in two to three sessions to examine functional connectivity in the brain. Each resting scan consisted of 36 consecutive base blocks, and participants were instructed to keep their eyes on the central fixation.

### Fixation-break Classification

To classify fixation-break vs. fixation-hold labels, support vector classification (SVC) was performed using leave-one-session-out cross-validation. The cross-validation procedure involved pre-learning, post-learning, and memory sessions. Participants performed a fixation-break task (2 runs, 90 TRs each) inside the scanner, where they were required to saccade to a new dot point in each TR that represented the break of fixation. The locations of dots matched the locations in which fractals were presented in the passive-viewing task. Data from both the fixation-break and resting-state tasks were used to train and validate the classifier, with an average validation accuracy of 99%. The best classifier was then used to predict fixation breaks in each TR of the fMRI sessions (pre-learning, post-learning, and memory). On average, the classifier predicted that fixation breaks occurred 27.3% of the time during these sessions. Notably, there were no significant differences in the number of fixation breaks observed in the base vs. probe (t12 = -0.10, p = 0.91), good vs. bad (t12 = 0.04, p = 0.96), and left vs. right blocks (t12 = -0.09, p = 0.92). These findings suggest that the classifier accurately predicted the occurrence of fixation breaks during the task and that the occurrence of fixation breaks was consistent across different experimental conditions.

### fMRI Data Acquisition

Functional and structural imaging was conducted on a 3 T Siemens MAGNETOM Prisma scanner equipped with a 64-channel head coil at National Brain Mapping Laboratory (NBML; Tehran, Iran). For each participant, a whole-brain structural image was acquired using a T1-weighted MPRAGE pulse sequence (TR = 1.8 s, TE = 3.53 ms, voxel size = 1 × 1 × 1 mm^3^, flip angle = 7^◦^). Functional images were collected using an EPI sequence (TR = 2 s, TE = 30 ms, voxel size = 3 × 3 × 3 mm^3^, flip angle = 90^◦^).

### Behavioral Analysis

Behavioral analyses were conducted in MATLAB (MathWorks). To determine whether the learning procedure differentially affected the performance and response time for the good and bad objects, a repeated-measures ANOVA was performed, followed by a Tukey multiple comparison test. Furthermore, average performance and reaction time across training and memory sessions were compared using t-tests.

### fMRI Data Analysis

fMRI data were analyzed using the AFNI/SUMA software package (Cox, 1996; Saad et al., 2004) and MATLAB (MathWorks) scripts. The pre-processing steps included: 1. Detrending and despiking (3dToutcount, 3dDespike); 2. Slice time correction (3dTshift); 3. Alignment to the anatomical T1 image (align_epi_anat.py); 4. Skull stripping and non-linear warping of the T1 image into MNI space (auto_warp.py); 5. Motion correction (3dvolreg) and 6. Non-linear warping of EPI data into MNI space (3dNwarpApply). The resulting EPI time series from all runs were concatenated and converted into percentage change from the mean. A general linear model (GLM) analysis was then performed to estimate the contribution of each factor to the percentage change from the mean (3dDeconvolve). The model consisted of six regressors ([good, bad] × [left, center, right]) convolved with the canonical hemodynamic response function. Six rigid-body motion parameters (x, y, z dimensions, roll, pitch, and yaw) were included in the GLM as additional regressors to account for the residual effects of participants’ motion. These regressors were entered into contrasts to yield “Good vs. Bad”, “Left vs. Right”, and “Center vs. Periphery” responses. After spatial smoothing with a 6 mm FWHM kernel (3dBlurInMask), linear contrasts were taken to a group-level analysis to calculate the mean and test whether the average estimates were significant (3dttest++). Finally, the cluster correction method was applied to find the significant clusters. Estimates of spatial smoothness based on the group residuals were found by simulating noise-only volumes, assuming the autocorrelation function (ACF) was given by a mixed-model, and then determining the threshold for the significant clusters (3dClustSim). The minimum cluster size for two-sided α < 0.05 and voxelwise significance of P < 0.05 was 127 and 134 voxels in the post-learning and memory sessions, respectively.

### Functional Connectivity Analysis

Resting-state correlation analysis was carried out using the residuals extracted from each region of interest (ROI). The anatomical parts that were significantly activated in the post-learning or memory session were included in each ROI. The physiological fluctuations of the white matter, the ventricles, and the cerebrospinal fluid (CSF) were regressed out of the residual time series, and bandpass filtering between 0.01 and 0.1 Hz was applied. Then Pearson’s correlation between the average signal of each pair of ROIs was calculated. The dissimilarity was measured as 1-ρ, with ρ being the pairwise Pearson’s correlation. Next, we applied multidimensional scaling to the dissimilarity matrix to visualize the resting-state distances in 2D (mdscale in MATLAB). Clustering of value coding ROIs based on their resting-state distance was implemented using the density-based clustering algorithm (DBSCAN). It required no prior knowledge of the number of clusters and allowed arbitrarily shaped clusters. The minimum number of data points to define a cluster was set to 2, and the radius of the neighborhood was set to the 75th percentile of all pairwise dissimilarities.

---

**Fig. S1.**
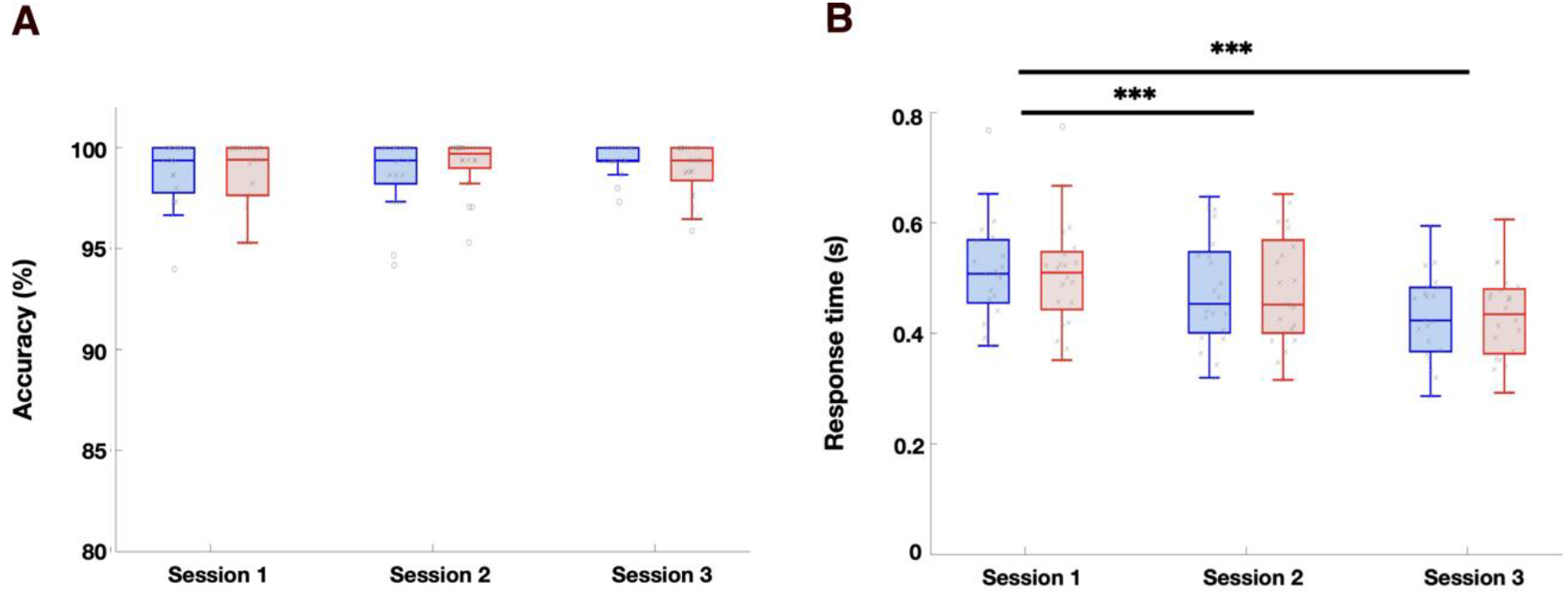
Behavioral results in force trials. (A) Performance in force trials across the three training sessions. Boxplots show the choice rate of good and bad objects in each session. The central mark within each box indicates the median, and the bottom and top of the box correspond to the 25th and 75th percentiles, respectively. The whiskers extend to the extreme values within 1.5 times the interquartile range. Outliers are plotted as dots. (B) Response time in force trials across the three training sessions. Same conventions as in (A) (*p < 0.05, **p < 0.01, ***p < 0.005).

**Fig. S2.**
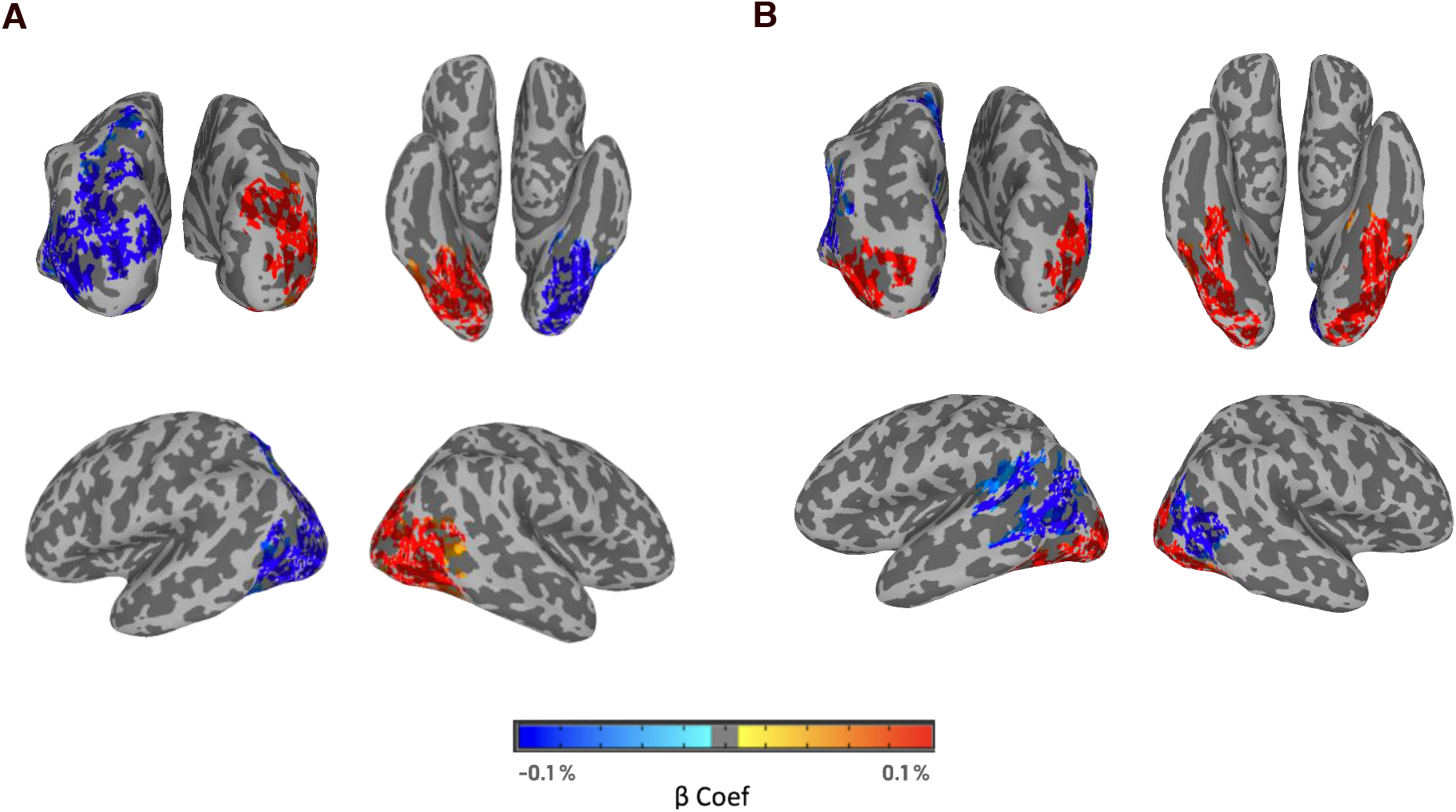
Brain regions with significant spatial coding. (A) Brain regions with significant hemifield selectivity. Warm colors show regions more strongly activated in response to the objects displayed on the left hemifield, and cool colors show the opposite (p < 0.05, α < 0.05 cluster corrected). (B) Brain regions with significant central vs. peripheral discrimination. Warm colors show areas more strongly activated in response to the objects displayed in the center, and cool colors show the opposite (p < 0.05, α < 0.05 cluster corrected).

**Fig. S3.**
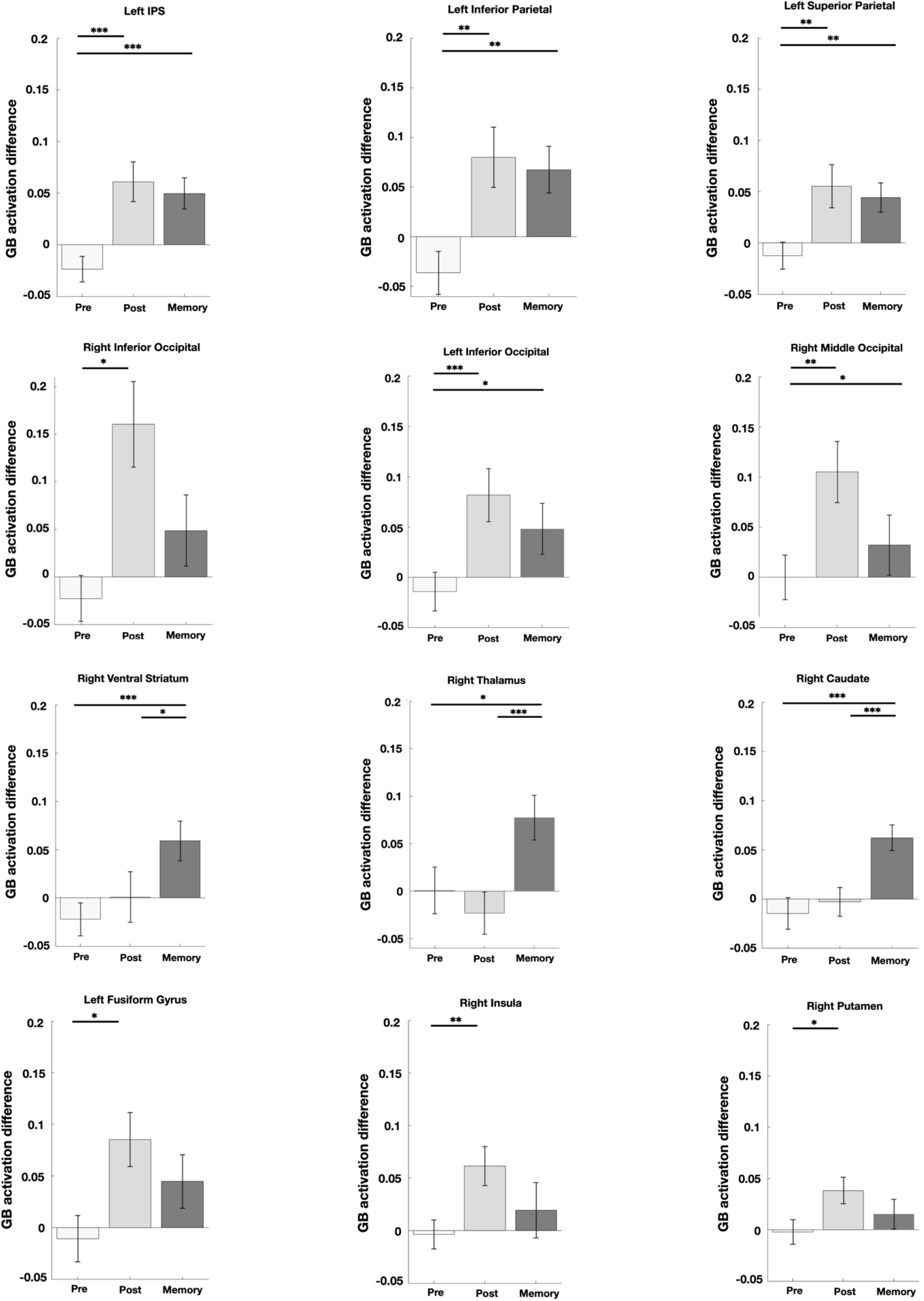
Difference in the response magnitude to good and bad objects in regions with significant activation in the post-learning or the memory session. Bar plots show the activation difference in the pre-learning, post-learning, and memory sessions (*p < 0.05, **p < 0.01, ***p < 0.005).

**Fig. S4.**
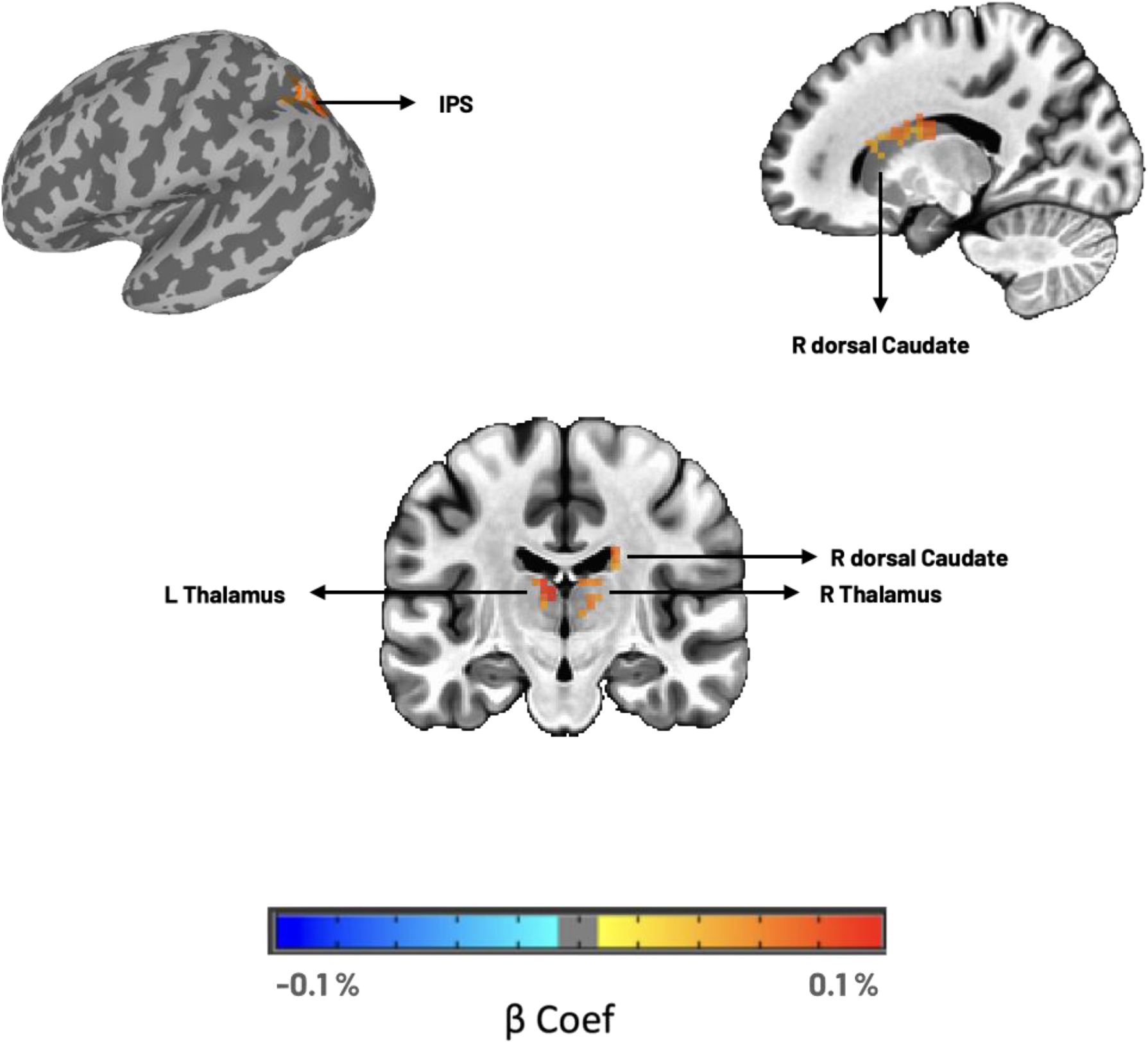
Regions showing significant activation in response to the memorized value. Brain regions with significantly different responses to the memorized good vs. memorized bad objects in the memory session (p < 0.05, α < 0.05 cluster corrected).

**Table S1.**
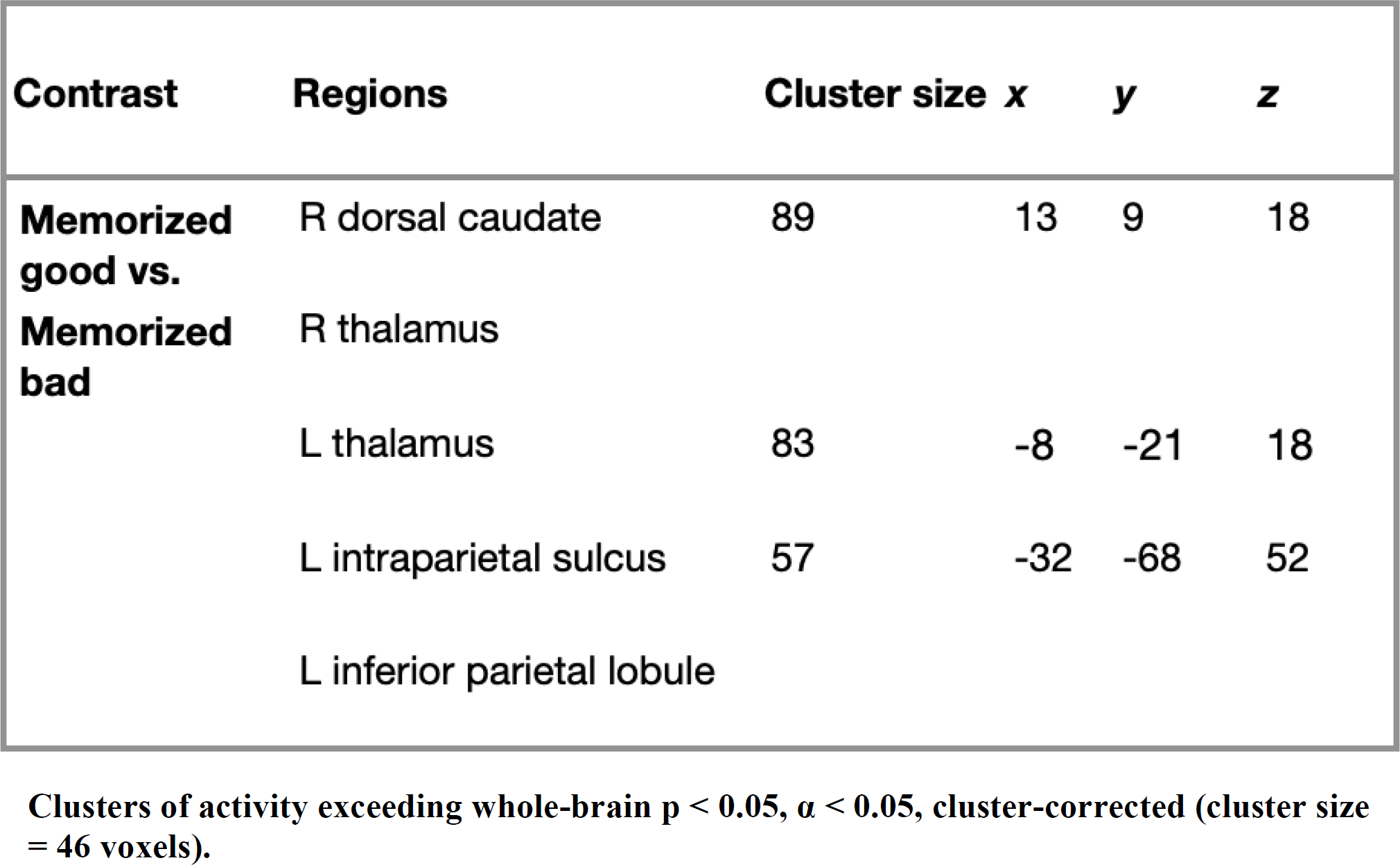
Summary of whole-brain analysis results.

| **Contrast** | **Regions** | **Cluster size** | **x** | **y** | **z** |
| --- | --- | --- | --- | --- | --- |
| **Memorized good vs. Memorized bad** | R dorsal caudate | 89 | 13 | 9 | 18 |
|  | R thalamus |  |  |  |  |
|  | L thalamus | 83 | -8 | -21 | 18 |
|  | L intraparietal sulcus | 57 | -32 | -68 | 52 |
|  | L inferior parietal lobule |  |  |  |  |

Clusters of activity exceeding whole-brain $p < 0.05$ , $\alpha < 0.05$ , cluster-corrected (cluster size = 46 voxels).

**Fig. S5.**
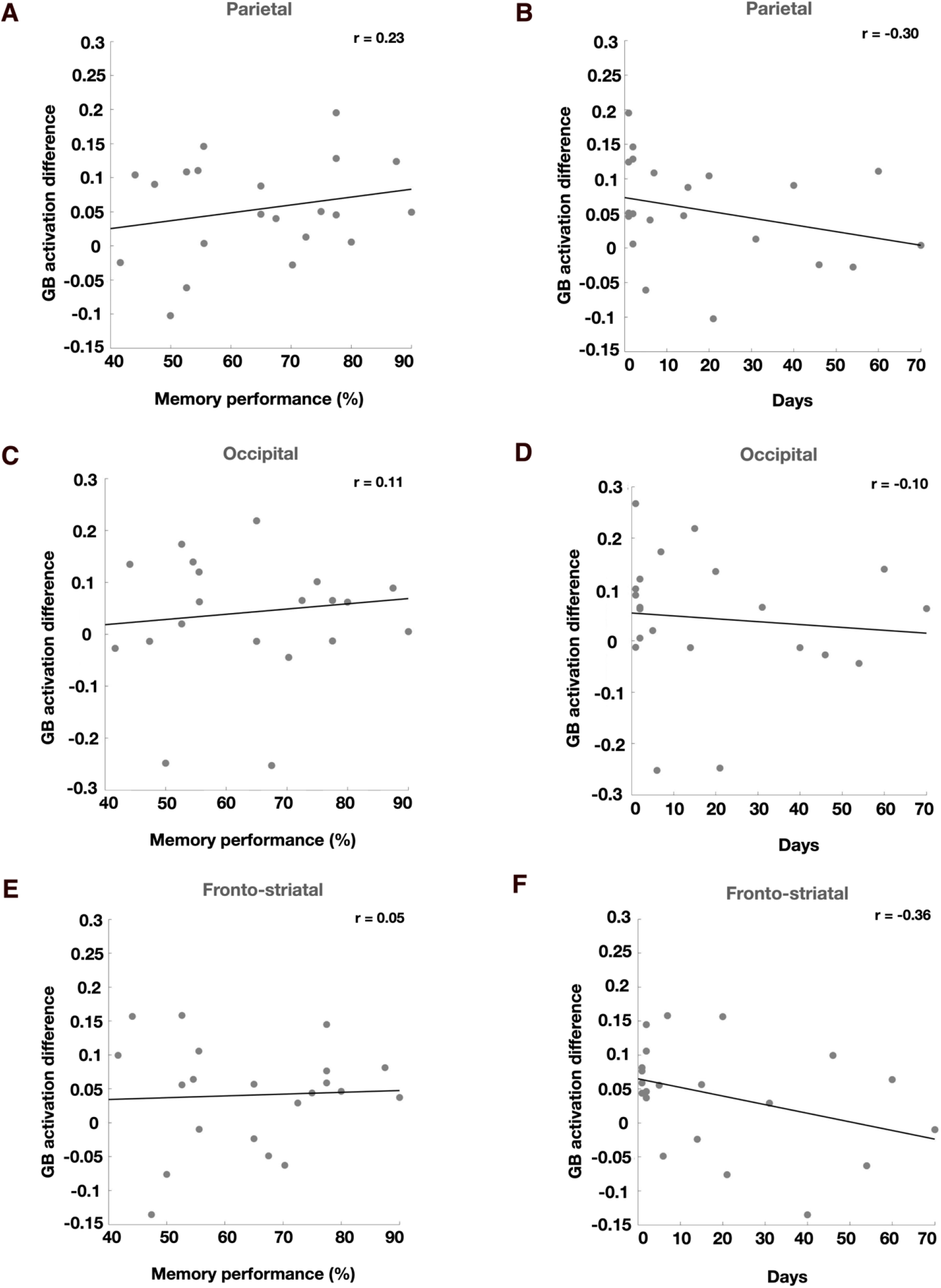
Relationship between brain activation and the days-long value test performance. (A) Same format as Fig. 5A, but for the parietal cluster. (B) Same format as Fig. 5B, but for the parietal cluster. (C) Same format as Fig. 5A, but for the occipital cluster. (D) Same format as Fig. 5B, but for the occipital cluster. (E) Same format as Fig. 5A, but for the fronto-striatal cluster. (F) Same format as Fig. 5B, but for the fronto-striatal cluster. (Pearson’s correlation “r” is noted in the plots; *p < 0.05, **p < 0.01, ***p < 0.005).

